# *Giardia* alters commensal microbial diversity throughout the murine gut

**DOI:** 10.1101/087122

**Authors:** NR Barash, JG Maloney, SM Singer, SC Dawson

## Abstract

*Giardia lamblia* is the most frequently identified protozoan cause of intestinal infection. Over one billion people are estimated to have acute or chronic giardiasis, with infection rates approaching 90% in endemic areas. Despite its significance in global health, the mechanisms of pathogenesis associated with giardiasis remain unclear as the parasite neither produces a known toxin nor induces a robust inflammatory response. *Giardia* colonization and proliferation in the small intestine of the host may, however, disrupt the ecological homeostasis of gastrointestinal commensal microbes and contribute to diarrheal disease associated with giardiasis. To evaluate the impact of *Giardia* infection on the host microbiota, we use culture-independent methods to quantify shifts in the diversity of commensal microbes throughout the entire gastrointestinal tract in mice infected with *Giardia*. We discovered that *Giardia’s* colonization of the small intestine causes a systemic dysbiosis of aerobic and anaerobic bacterial taxa. Specifically, giardiasis is typified by both expansions in aerobic *Proteobacteria* and decreases in anaerobic *Firmicutes* and *Melainabacteria* in the murine foregut and hindgut. Based on these shifts, we created a quantitative index of murine *Giardia*-induced microbial dysbiosis. This index increased at all gut regions during the duration of infection, including both the proximal small intestine and the colon. Thus giardiasis could be an ecological disease, and the observed dysbiosis may be mediated directly via the parasite’s unique anaerobic fermentative metabolism or indirectly via parasite induction of gut inflammation. This systemic alteration of murine gut commensal diversity may be the cause or the consequence of inflammatory and metabolic changes throughout the gut. Shifts in the commensal microbiota may explain observed variation in giardiasis between hosts with respect to host pathology, degree of parasite colonization, infection initiation, and eventual clearance.

## INTRODUCTION

*Giardia lamblia* is a microaerophilic protozoan parasite of humans and animals causing significant morbidity and diarrheal disease worldwide (1, 2). Giardiasis is a zoonotic disease with diverse animal reservoirs. Parasites infect and complete their life cycle in mammalian hosts, and infected animals shed *Giardia* cysts into water supplies. Over one billion people are estimated to have acute or chronic giardiasis, and rates of giardiasis approach 90% in endemic areas (3, 4). When prevalent, giardiasis has been implicated as a primary cause of growth restriction for children, resulting in long-term consequences such as stunting, failure to thrive, malnutrition, and cognitive disabilities (1, 2, 5). In addition, *Giardia* has been associated with substantial post-clearance irritable bowel symptoms in both children and adults (1, 6-8). The significant and adverse impact of giardiasis on global human health contrasts with a considerable lack of research efforts targeting prevention and treatment (9).

Common symptoms of acute and chronic giardiasis include abdominal cramps, gas, nausea, and weight loss. Severe giardiasis results in malabsorptive diarrhea with fatty, bulky stools (10). At the cellular level, parasite colonization of the gut is associated with malabsorption of glucose, salts, and water, disruption of intestinal barrier function, induction of enterocyte apoptosis, inhibition of brush-border disaccharidases and lactases, loss of mucosal surface due to increased crypt:villus ratios and shortened microvilli, and manipulation of host immune responses via arginine/nitric oxide limitation (1, 11-14).

Despite its global importance as a primary cause of diarrheal disease, how *Giardia* actually induces giardiasis remains enigmatic. *Giardia* produces no known toxin, and parasite colonization does not induce a robust inflammatory reaction (4, 15). Symptoms are generally believed to be the consequence of host tissue damage caused by the direct contact of the parasite with the intestinal villi (12). Paradoxically, giardiasis can frequently be asymptomatic (1) and parasite colonization can be associated with histologically normal tissue (16). The precise mechanism by which *Giardia* colonization and proliferation in the gastrointestinal tract induces diarrheal disease remains unclear (4).

An ecological perspective on giardiasis could help illuminate our understanding of *Giardia’s* pathogenesis (17). Mammalian gut commensal microbes are organized into specific trophic structures and benefit the host by breaking down complex substrates, supplying essential nutrients, and defending against opportunistic pathogens (1). Commensal microbes from all three domains of life colonize the gastrointestinal tract of mammals, with approximately 10^9^ cells /g in the small intestine and 10^12^ cells/g of luminal contents in the large intestine (1). The maintenance of these diverse types of gut microbes is critical for overall stability of this ecosystem (18). Microbial diversity can confer resilience and flexibility of community responses to various external stresses, both promoting and modulating the development of the immune system. Commensal gut microbial communities are known to both limit and exacerbate pathogen colonization (19-22), and the disruption of the gut ecosystem may impact *Giardia* colonization and associated disease symptoms. The histopathology of giardiasis has been noted to be less severe in germ-free mice than in conventional mice, suggesting that the presence of the intestinal microbiota can aggravate parasite infection or symptoms (23).

*Giardia* colonization of the mammalian host gut can be viewed within the context of invasion biology, in which an exotic invading species has cascading effects on the resident community (24). In conjunction with direct effects on host tissues, *Giardia* could cause disruptions to the ecological health of commensal microbes in the gastrointestinal tract, resulting in disease symptoms associated with giardiasis. *Giardia* colonization may alter microbial composition, metabolic capacities, or chemical homeostasis in the gastrointestinal lumen. Other gut ecologically-mediated diseases include infectious diseases with known bacterial pathogens (e.g., *Clostridium difficile or Salmonella typhimurium*) and immune-mediated diseases wherein the entirety of the gut community appears perturbed in the absence of a dominant pathogenic species (e.g., Crohn’s disease and irritable bowel disease (IBD)) (22, 25-28). Similarly, the eukaryotic apicomplexan parasite *Toxoplasma gondii* is proposed to act as a molecular adjuvant to the small intestinal microbiota, inducing acute ileitis (29). *T. gondii* infection induces a rapid overgrowth of Proteobacteria within the small intestine, aggravating immunopathology and promoting an anti-parasitic immune response (30, 31).

To evaluate potential dysregulation of commensal gut microbial ecology associated with giardiasis, we infected mice with *Giardia* and used standard cultivation-independent methods (32-37) to describe and quantify shifts in microbial diversity throughout the entire gastrointestinal tract. We demonstrate that *Giardia’s* colonization of the small intestine causes a systemic imbalance (dysbiosis) throughout the murine gut that persists during infection. We propose that *Giardia* infection disrupts commensal gut microbial ecology and metabolism in the host. This dysbiosis may be mediated directly via the parasite’s unique anaerobic metabolism and/or indirectly via parasite induction of host responses. This systemic alteration of murine gut ecology may be the cause or consequence of micro-inflammatory and metabolic changes throughout the gut, and could also explain observed variation in giardiasis with respect to host pathology, degree of parasite colonization, infection initiation, and eventual clearance.

## Materials and Methods

### Giardia growth conditions and infections

*Giardia lamblia* strain GS/M/H7 (ATCC 50801) was cultured in modified TYI-S-33 medium supplemented with bovine bile and 5% adult and 5% fetal bovine serum [56] in sterile 13 ml screw-capped disposable tubes (BD Falcon) incubated upright at 37°C without shaking.

For infections, eight week old C57BL/6J female mice (Jackson Laboratories) were kept under specific-pathogen-free conditions. Mice were kept as cage mates, with four mice per cage. The experimental design included two cohorts of sibling mice (n=32). Prior to inoculation, one cohort of sixteen mice was treated for four days with antibiotics (0.5 mg/ml vancomycin, 1 mg/ml neomycin, 1 mg/ml ampicillin) *ad libitum* in their drinking water to promote parasite colonization (38). Antibiotics were maintained in drinking water throughout the infection. Sixteen control mice were not given antibiotics. Study animals in each cohort were gavaged with one million *Giardia lamblia* GS (n=12) trophozoites or a saline vehicle control (n=4 per antibiotic group). Mice in replicates of four were sacrificed at three, seven and fourteen days post inoculation. Infection dates were staggered such that all sacrifices occurred within a three-day window. All animal studies were performed with IACUC approval at Georgetown University (Steven Singer, PI).

### Anatomical sampling of the murine gastrointestinal tract for microbial community analyses

For comparisons of microbial diversity in various regions of the murine gut during giardiasis, uninfected and *Giardia* infected mice were sacrificed at three, seven and fourteen after inoculation with *Giardia*. Anatomical sampling of the gut yielded six community DNA samples: mucosal and luminal proximal small intestine, mucosal and luminal distal small intestine, cecal contents, and colonic contents (See Figure 2). The proximal small intestine (pSI) was sampled as a 3 cm segment 3 cm distal to the pylorus, and the distal small intestine (dSI) was sampled as a 3 cm segment 10 cm distal to the pylorus. For the proximal and distal small intestinal samples, the luminal and mucosal contents were extracted separately. Luminal contents were isolated by gentle pressure, and the mucosa was removed by gentle scraping with a sterile glass slide. The cecum was opened with a sterile blade and 200 μl of cecal contents were removed. Colonic contents (COL) were sampled by excising the colon and removing two fecal pellets. Pelleted colonic samples from cage-mate mice exposed to the same diet were similar in weight and size. No samples were taken from the stomach. All samples were immediately flash frozen in liquid nitrogen and stored at -80C for subsequent microbial diversity analyses.

### Total gut microbial community DNA extraction

Total community DNA was extracted from 192 gastrointestinal tract samples from the 32 mice using standard methods with mechanical disruption (35) for paired-end sequencing with the llumina MiSeq. Small subunit rDNA primers (515F and 806R) were used to amplify total community DNA was according to protocols developed at the Earth Microbiome Project (36, 37). Small subunit rDNA amplicon sequencing was performed at Argonne National Laboratories using protocols from the Earth Microbiome Project (37).

### Amplicon assembly and identification of OTUs

More than two million total reads were used for analysis after sequence quality filtering and assembly of paired end reads. Default parameters were used to pair forward and reverse reads with SeqPrep (https://github.com/jstjohn/SeqPrep) and call OTUs with open-reference picking at 97% confidence using the GreenGenes 13_8 database. Singleton OTUs were discarded. Analyses of alpha and beta diversities were performed in QIIME pipeline (39) using default parameters for OTU tables rarefied to 3000 sequences/sample. OTU tables were filtered to reflect only antibiotic treated, antibiotic naïve, foregut, hindgut, or any other subclassifications as needed for diversity analyses. Samples with less than 3000 reads were discarded.

### Analyses of gut microbiome diversity

All microbial diversity metrics were calculated using rarefied OTU tables at 3000 sequences/sample, with the exception of DESeq (used for unrarefied OTU tables with a 3000 sequences/sample minimum count) (40). Alpha diversity for each sample was evaluated using observed OTUs, Chao1, Shannon, and whole tree Phylogenetic Distances. PCOA plots were visualized using EMPeror (41) and both unweighted and weighted UniFrac were used for further analyses (42, 43). ADONIS analysis was performed on unweighted and weighted Unifrac matrices to calculate P values. All diversity metrics were calculated using weighted UniFrac. Similar trends were seen using both weighted and unweighted algorithms (unweighted R2 values presented in Supplemental Table 2). Mucosal and luminal samples from the same small intestinal site were combined unless otherwise indicated. For overall summaries of microbial diversity, proximal and distal small intestine sample were combined and designated “foregut”, whereas combined cecum and colon samples were designated “hindgut”.

Taxa with significant variation (e.g., dynamic taxa) were identified between anatomical sites in experimental treatments using a combination of group_significance.py script to perform a Kruskal-Wallis test on rarefied OTU tables and differential_abundance.py on unrarefied OTU tables to implement the DESeq algorithm (40, 44). Giardiasis Indicator taxa were defined as taxonomic groupings identified through both DESeq and Kruskal-Wallis analyses that had consistent variation in diversity (corrected P-values <0.05), and fold change in relative read abundance in infected animals was calculated relative to relative read abundance in uninfected animals. Lastly, the Murine Giardiasis Dysbiosis Index (MGDI) was calculated using calculate_taxonomy_ratios.py in QIIME, specifying increased *Moraxellaceae* and *Rhodocyclaceae* with decreased *Clostridiales*. PCOA plots with explicit axes were created using the Make_emporer.py script in QIIME.

### Quantitation of parasite density (burden) using QPCR

For the QPCR based analysis of microbial abundance, the gastrointestinal tracts of euthanized mice were isolated and frozen as described above. Specifically, a 3 cm segment located 7 cm distal to the pylorus (proximal small intestine) was used in its entirety for *Giardia* quantification. QPCR of an intragenic region unique to Chromosome 1 was performed with *Giardia*-specific primers GS-1-318F GCAGAAACAGTGCTTTGAGG and GS-1-318R TTGTTTACGGCAAGGAAATG. DNA was extracted using phenol-chloroform and bead beating (45). The single copy, stably expressed murine nidogen-1 (*nid1*) gene was used as an internal control to quantify the contribution of murine DNA to intestinal segments with primers nido.F CCAGCCACAGAATACCATCC and nido.R GGACATACTCTGCTGCCATC (46). The differential counts to threshold (ΔCT) between *nid1* and *GS-1-318* was calculated and ΔCT normalized against murine contribution. Standard curves for QPCR were created using known concentrations of *Giardia* GS DNA. The averaged CT of three technical replicates was utilized to extrapolate the concentration of *Giardia* per sample. The fold change was calculated between averaged negative samples and each infected sample.

## Results

### Giardia colonization is correlated with alterations in host commensal diversity in both antibiotic-treated and antibiotic-naive mice

To quantify how *Giardia* infection alters the diversity and abundance of host-associated gut microbiota, we compared the gut microbiota of mice infected with *Giardia* to the microbiota of uninfected mice gavaged with a saline vehicle control (Methods). Antibiotic treatment is generally required for robust experimental *Giardia* infections in mice (47, 48); To investigate potential colonization resistance, we compared infections and commensal diversity in both antibiotic-naïve and antibiotic-treated animals. *Giardia lamblia* strain GS (Assemblage B) was used for all infections, as this strain colonizes antibiotic-free mice more readily than the Assemblage A strain WBC6 (38) (47, 49) (Figure 1). A total of 192 samples from the intestinal tract of 32 animals in 8 treatment groups, sampled from 6 sites (luminal and mucosal sampling of proximal and distal small intestine, bulk sampling of cecum and colon) were used for diversity analyses.

**Figure 1.**
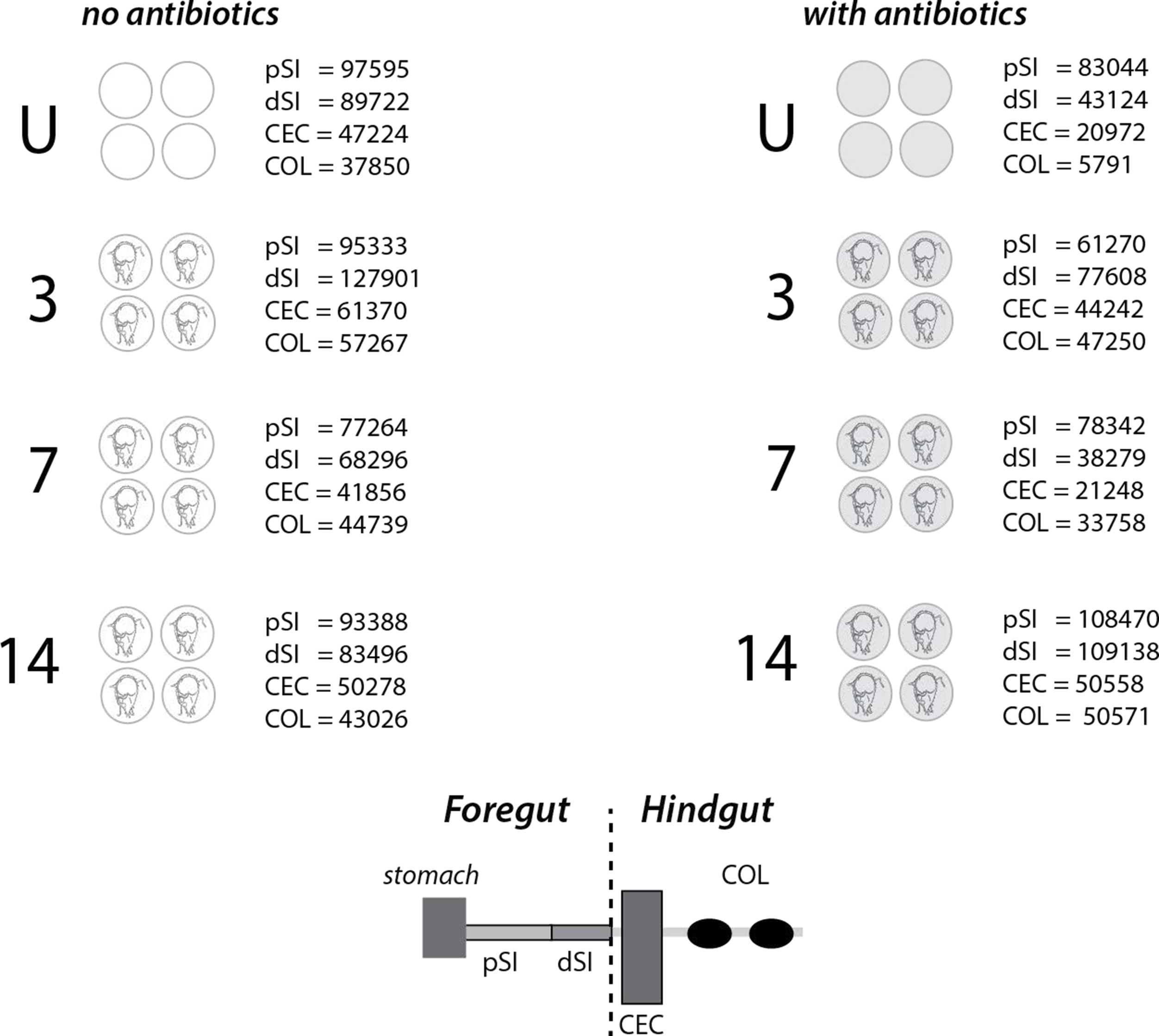
Sampling scheme for querying murine gut ecological shifts during giardiasis with and without antibiotic treatment. Two cohorts (N=16) of sibling mice – one not treated with antibiotics, and one treated with antibiotics (gray, see Methods)—were used to test the potential effects of *Giardia* colonization on the microbial diversity of the murine gut. Study animals were infected with *Giardia lamblia* GS trophozoites (N=12 animals per treatment group), or a saline vehicle control (Uninfected (U), with N=4 per treatment group) by gavage. Cohorts of infected animals were sacrificed at 3, 7 and 14 days post-gavage (N=4 per day). Gut intestinal tracts from each treatment group and day post-inoculation were dissected for subsequent diversity analyses (proximal small intestine (pSI), distal small intestine (dSI, cecum contents (CEC) and colonic contents (COL). The total number of reads per cohort (N =4) are shown for each experimental condition. Luminal and mucosal reads were pooled for most analyses (see Methods). The total reads per animal within each cohort in each experimental condition are summarized in Supplemental Table 1.

**Figure 2.**
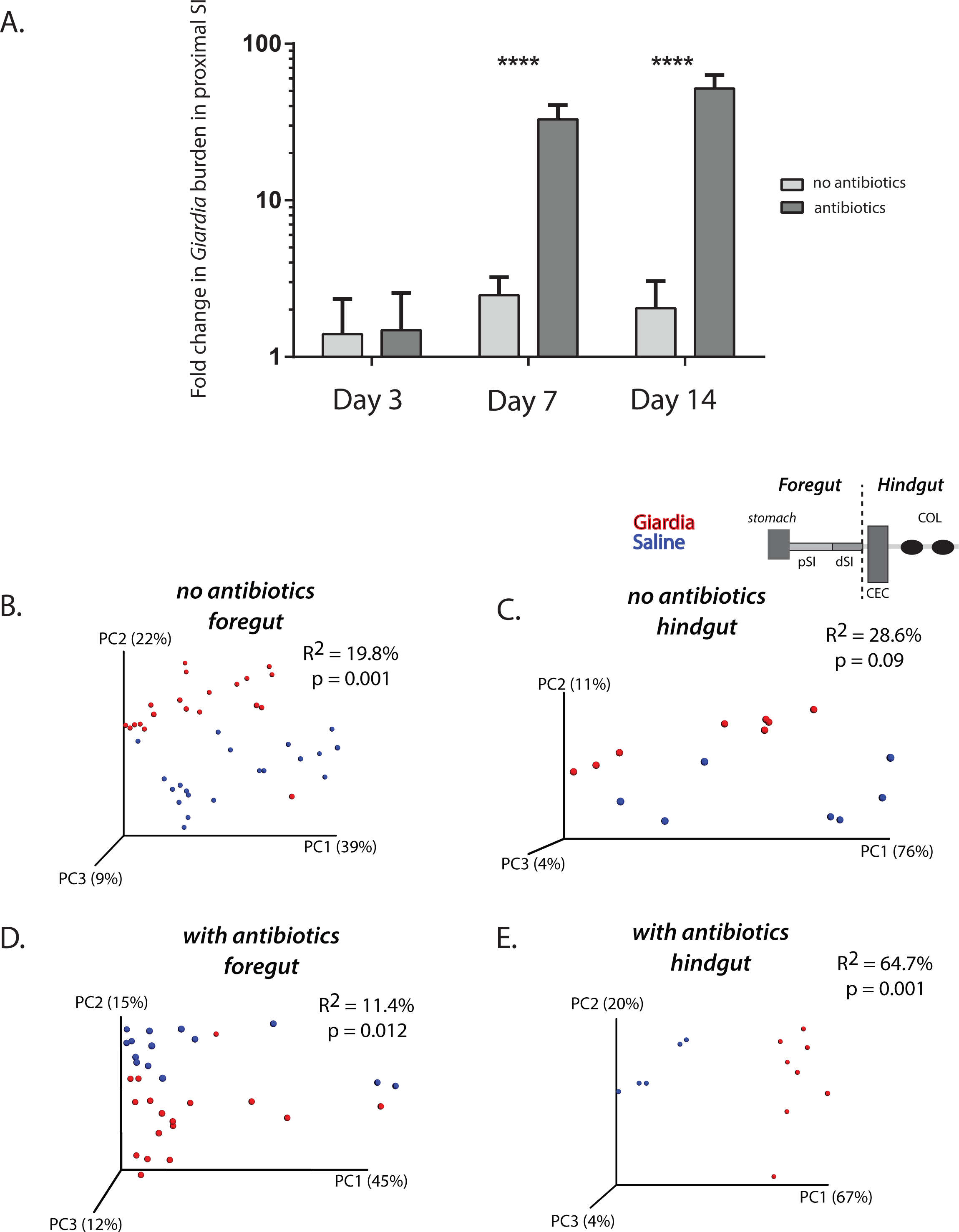
Giardia infection significantly alters foregut and hindgut microbial diversity in antibiotic treated and naïve mice. The comparison of the fold change in *Giardia* burden in proximal small intestinal samples (pSI) by QPCR between antibiotic-treated and non-treated animals is presented at days 3, 7 and 14 post *Giardia* gavage (A). Asterisks indicate significant differences as assessed by Sidek’s Multiple Comparison Test. In B-E, weighted Unifrac comparisons of beta diversity are shown using Principal Coordinate plots (PCOA) comparing the variance in the diversity of all foregut samples and hindgut samples with and without antibiotic-treatment. Samples with a saline vehicle control are shown in blue, and samples after two weeks of *Giardia* infection are shown in red (B-E). Significance (p) and strength of explained variation (R2) were assessed with ADONIS.

For diversity analyses, total community DNA was extracted from each sample and amplified using universal small subunit (ssu) rDNA primers (37). Amplified ssu genes were sequenced using high-throughput methods (see Methods) to generate 2,002,493 paired-end Illumina reads. A total of 12,171 individual OTUs were called at the 97% identity threshold from all samples (Figure 1 and Supplemental Table 1). The total numbers of sequence reads for each anatomical site ranged from about 6000 to over 60,000 (Figure 1). Both antibiotic-treated and naïve mice had a median unique OTU count of 468.

To quantify the degree of parasite infection (or *Giardia* burden) in the antibiotic-treated and untreated cohorts, we used *Giardia*-specific quantitative PCR of a single-copy intragenic region (GS-1-318). Proximal small intestinal samples (3 cm segments located 7 cm distal to the pylorus) were collected from animals in both treatment groups on day 3, day 7 and day 14, and the *Giardia* burden was assessed using QPCR. Without antibiotic pretreatment, we noted moderate infection density that peaked on Day 7 (Figure 2A). With antibiotic pretreatment, however, we saw a significant 80-fold increase in *Giardia* abundance relative to untreated mice on Day 7, and a further increase in parasite abundance by Day 14 post-infection, when *Giardia* burden in untreated mice had declined.

To investigate potential colonization resistance in the murine giardiasis model, we infected both antibiotic-treated and antibiotic naïve mice with *Giardia*. Although the overall parasite burden was lower without antibiotic treatment (Figure 2), colonization was achieved and the microbiota varied similarly to antibiotic-treated, *Giardia*-infected mice. Specifically, with respect to overall microbial diversity between all sequences in the foregut and hindgut (Figure 2B-E), we confirmed that *Giardia* infection results in significant differences in the variances of foregut and hindgut sequences, in both non-treated and antibiotic treated animals.

### Overall species richness and evenness are not significantly altered within each anatomical site during giardiasis

To evaluate the overall dynamics of ecological shifts in gut microbiota during giardiasis, we first analyzed species richness and evenness (alpha diversity) using QIIME (Figure 3 and Supplemental Figure 1). Sampling accuracy for giardiasis samples with antibiotic treatment (Figure 3A) and without (Supplemental Figure 1A) was evaluated for all anatomical samples at each time point post infection using rarefaction of Chao1 estimates (32-34). For both treatment groups, the Chao1 for all samples (all anatomical sites) by day was estimated to 3000 OTUs.

**Figure 3.**
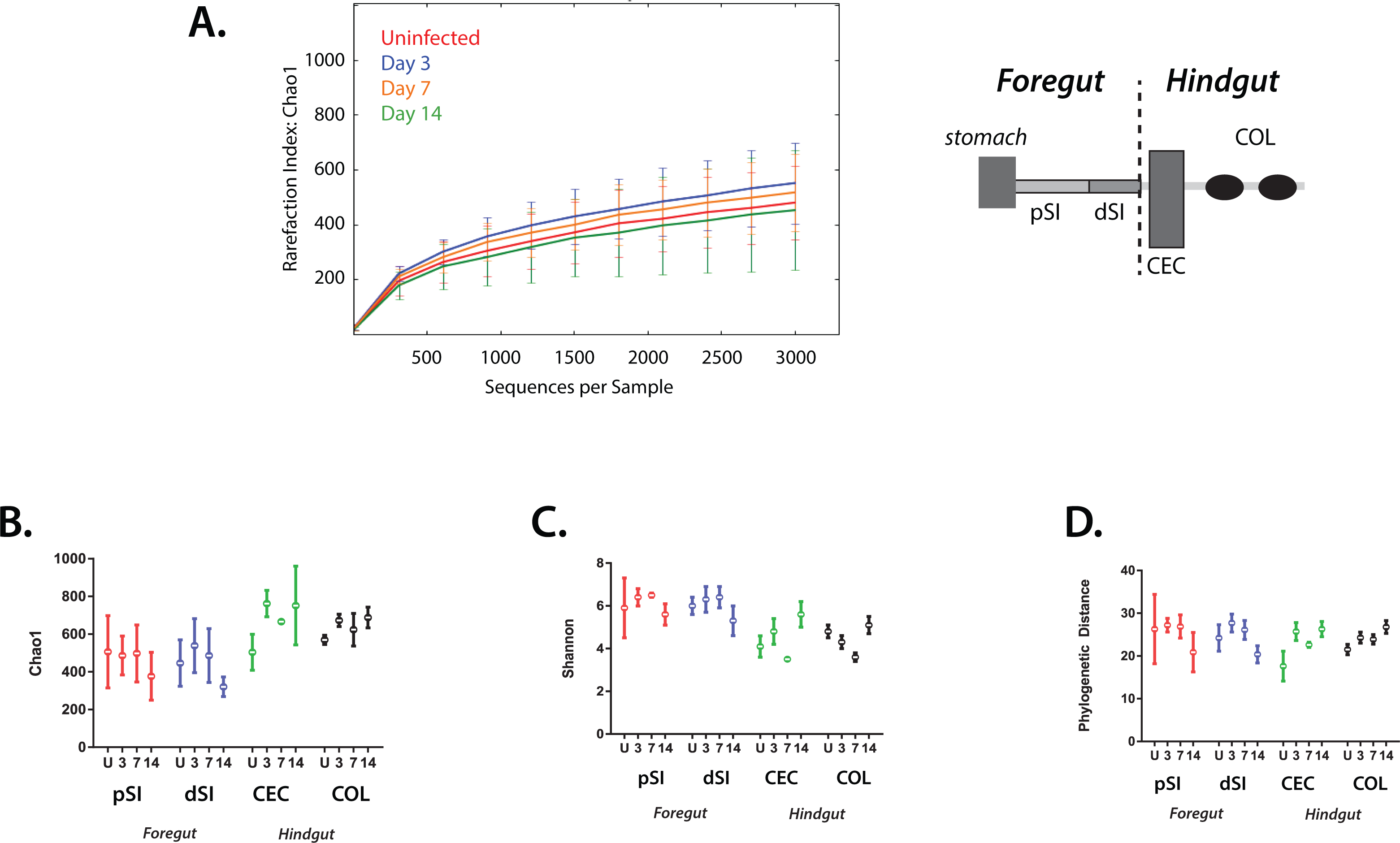
Overall alpha diversity by intestinal site is not significantly changed during giardiasis. Sampling adequacy (alpha diversity) was assessed using rarefaction plots (A) to compare all samples using the Chao1 estimate during the course of giardiasis (uninfected, day 3, day 7, and day 14) with antibiotic treatment (see Supplemental Figure 1 for no antibiotic treatment). In B-D, alpha diversity measures of species richness, evenness, and phylogenetic distance summarize microbial diversity in regions of the murine gut (proximal small intestine (pSI), distal small intestine (dSI, cecum contents (CEC) and colon (COL) in control, antibiotic treated, non-infected cohorts (U) and in infected cohorts (days 3, 7, and 14). For these analyses, the OTU tables were rarefied to 3000 reads/sample and analyzed in QIIME (see Methods) to compare the relative read abundances. Chao 1 estimates of species richness for both antibiotic treated are shown (B) where increased Chao1 values correspond to increased numbers of unique taxa. Species evenness was also estimated using the Shannon index (C) where a high Shannon value indicates equal distribution between species and a low Shannon value indicates dominance by certain taxa. Lastly, the Phylogenetic Distance (D) was used to estimate the evolutionary relationships of the community; a higher phylogenetic distance value corresponds to a more diverse community. Error bars correspond to variation between gut samples of each of the four animals in that experimental group.

To compare alpha diversity between sites, four pooled anatomical samples were compared: proximal and distal small intestine (pSI and dSI), cecum (CEC) and colon (COL) (Figure 3B-2D). Species richness was similar for all samples based on Chao1 and observed OTUs calculations. For any day of infection, regardless of antibiotic treatment, the proximal and distal small intestine samples were slightly less rich in unique OTUs than the cecal and colonic samples (Figure 3B and Supplemental Figure 1B). This trend was more pronounced in the antibiotic-naïve cohort (Supplemental Figure 1B).

To assess taxonomic evenness of the gut microbiota, we performed estimates of the Shannon Index for each experimental condition and anatomical site (Figure 3C and Supplemental Figure 1C). Higher Shannon Index values imply an even distribution of abundances, in contrast to low Shannon values that suggest certain community members predominate. We found that antibiotic treatment increased the Shannon Index in the proximal and distal small intestine compared to antibiotic-naïve mice, but did not affect the cecal or colonic samples. Over the time of infection, microbiota diversity at all gut sites remained even with respect to OTU abundances. During giardiasis, the antibiotic-treated cohort exhibited more fluctuation than the non-treated cohort (Figure 3C and Supplemental Figure 1C).

Lastly, we quantified phylogenetic distances between taxa during giardiasis in the four pooled anatomical samples (Figure 3D and Supplemental Figure 1D). For antibiotic treated mice, we observed that phylogenetic distances remained consistent over the time course of infection and between body sites. When individual antibiotic-treated mice were compared, the proximal and distal small intestines exhibited increased variability in phylogenetic distances, as indicated by wider error bars, as compared to the cecum and colon. Comparing phylogenetic distances between all gut sites and time points, we found that antibiotic treated mice had a higher estimate of phylogenetic distances between taxa than non-treated animals.

### Giardia infection is associated with a significant dysbiosis within the murine foregut and hindgut

While *Giardia* trophozoites preferentially colonize the most proximal small intestine (Figure 2A and (50-52)), they are also found in the distal small intestine, as well as the cecum and large intestine (50-52). Within the antibiotic-treated proximal small intestine, we found that *Giardia* infection substantially changed the diversity of the host microbiota during the fourteen-day course of infection (Supplemental Figure 2A). Although mucosal and luminal communities are generally distinct in the large intestine (53), luminal/mucosal proximal and distal intestinal samples were not significantly clustered with PCOA (Supplemental Figure 3B) and were pooled for subsequent taxa-specific analyses. Total cecum and colonic samples did not significantly cluster in PCOA during giardiasis (Supplemental Figure 3C) and were pooled for subsequent analysis.

We used weighted UniFrac analysis to evaluate whether giardiasis is a predominant contributor to changes in murine gut microbial diversity (Supplemental Figure 2). By comparing sequences from uninfected animals to those from animals fourteen days post-*Giardia* infection, we found that sequences from any small intestinal site clustered together based on giardiasis status, rather than with respect to small intestinal subsite (Supplemental Figure 2D).

Antibiotic treatment by itself changes the gut microbial ecosystem. Without antibiotics, we found that the proximal small intestinal microbiome was dominated by the *Proteobacteria* and *Firmicutes* bacterial phyla, and that antibiotic treatment supported a larger population of *Bacteroidetes* and *Melainabacteria* compared to naive individuals (Figure 4). While we noted these significant differences in antibiotic versus non-antibiotic treated samples, we highlight our subsequent analyses of the treatment group with antibiotics, as this group had significantly more *Giardia* abundance, particularly by day 14 post infection (Figure 2). Specifically, we focused our attention on comparative analyses of antibiotic-treated animals with and without giardiasis by anatomic site. All cognate analyses for animals not treated with antibiotics are presented in Supplemental Figures (Supplemental Figures S4-S7).

**Figure 4.**
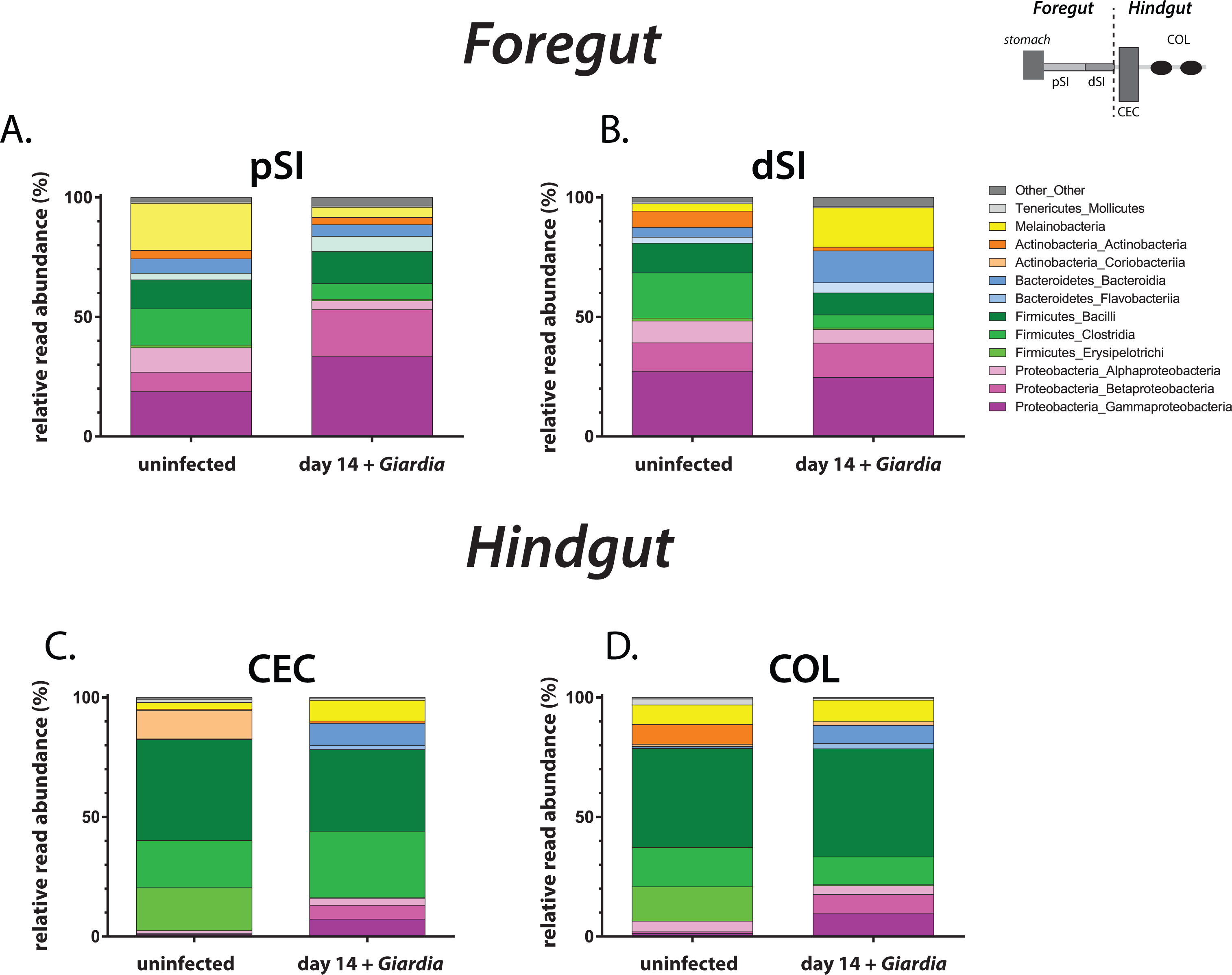
Significant changes in gut microbiome diversity in the proximal small intestine and colon during giardiasis. Fourteen days post infection, the relative read abundance (%) of the most abundant bacterial divisions and phyla are shown for the mouse foregut: (A) proximal small intestine (pSI), (B) distal small intestine (dSI), and the mouse hindgut: (C) cecum contents and (D) colon (COL). Each anatomic section is compared to the cognate uninfected sample.

### Increased Proteobacteria diversity and decreased Firmicute diversity exemplify the primary ecological changes during giardiasis

In the proximal small intestine (pSI) of antibiotic-treated animals, the diversity of several taxonomic groups was substantially altered during giardiasis (Figure 4A). Specifically, the *Proteobacteria* expanded from 33% to 56% of total reads after fourteens days of infection. The expansion of both beta- and gamma-*Proteobacteria* diversity accounted for over 80% of changes in unique *Proteobacteria* OTUs. In contrast, the overall percentage of *Firmicutes* reads decreased from 28% to 20% following infection. The significant decrease in *Clostridia* taxa during giardiasis accounted for over 80% of the changes in *Firmicutes* OTUs. Lastly, *Giardia* infection reduced diversity of *Melainabacteria* from almost 20% to 4% with respect to total reads in the proximal small intestine. We also saw alterations in the diversity of commensal microbes in the distal small intestine (dSI, Figure 4B), with increases in the diversity of both *Melainabacteria* and *Firmicutes*.

We observed similar trends in changing diversity in the foregut (pSI and dSI) within the antibiotic-naïve mice during giardiasis (Supplemental Figures 4 and 5). With respect to the proximal small intestine, the overall diversity of the *Proteobacteria* expanded from 14% to 21% of total reads, with the beta-*Proteobacteria* in particular increasing from just over 1% of reads to almost 8% during giardiasis. Infection induced a decrease in the Clostridia from 25% to 8% of total reads. The *Melainabacteria* decreased from 4% to 1% of total reads. Clustering by treatment is evident in PCOA plots of antibiotic treated and naïve mice, and ADONIS analysis showed a similar impact of *Giardia* infection, accounting for 13% of total variance in both antibiotic-treated and naïve groups (Supplemental Figure 5). In the distal small intestine (dSI, Supplemental Figure 4B), we observed slight decreases in the diversity of both *Melainabacteria* and *Firmicutes*.

Due to host peristalsis and inflammatory responses occurring along the entire gastrointestinal tract, we predicted parasite colonization of the foregut could induce dysbiosis in the hindgut, far from the site of infection. We discovered that hindgut samples also have significant overall changes in bacterial diversity following fourteen days of giardiasis in both antibiotic treated and non-treated animals (Figure 4 and Supplemental Figures S3-S5). During giardiasis, *Firmicutes* diversity in the cecum decreased as the *Proteobacteria* diversity increased. Similarly, large intestinal diversity was dominated by *Firmicutes* taxa, with an expansion of the *Proteobacteria* in antibiotic treated animals (Figure 4D). The *Melainabacteria* accounted for 8% of reads, and thus were unchanged during infection. During giardiasis without antibiotic treatment, both the cecum and large intestinal samples are composed predominantly of *Firmicutes* and *Bacteroidetes*, which together account for about 94% of reads regardless of infection (Supplemental Figure 4).

What are the primary bacterial taxa, or “indicator” taxa that shift in abundance during giardiasis? We used several multivariate statistical methods (DESeq2 and ANOVA; QIIME 1.9) to query microbial taxa that significantly changed after two weeks of giardiasis (40, 44). The OTUs with significant variation in relative abundance during infection ranged between 20-50% of total reads, depending on the day, site, and condition analyzed. Overall, *Clostridiae* and *Proteobacteraciae* are the taxa with the most dramatic changes in diversity throughout both the foregut and hindgut during giardiasis (Figure 5).

**Figure 5.**
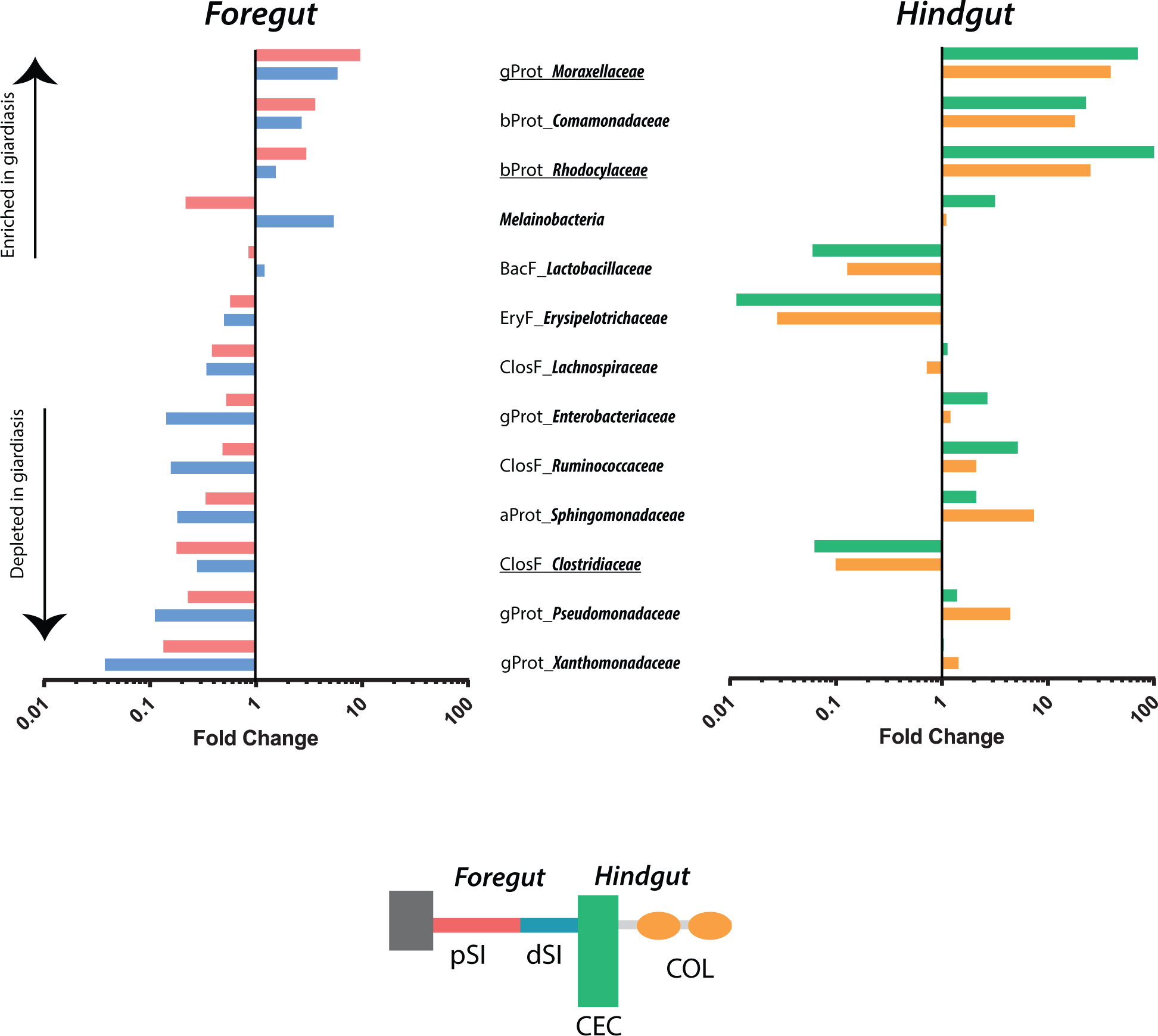
Significant shifts in beta diversity in both the murine foregut and hindgut during giardiasis. The relative abundance of bacterial taxa that were significantly enriched or depleted after fourteen days of *Giardia* infection is plotted by comparing the fold change in uninfected to infected animals using the same anatomical sample. The taxonomy of each species is indicated, where aProt = alpha-Proteobacteria, bProt = beta-Proteobacteria, gProt = gamma-Proteobacteria, BacF = Bacilli (Firmicutes), EryF = Erysipelotrichi (Firmicutes), ClosF = Clostridia (Firmicutes). Melainabacteria were categorized as Cyanobacteria-Chloroplast in QIIME. Underlined taxa are used to calculate the Murine Giardiasis Dysbiosis Index (see Figure 7 and Supplemental Figure 7).

To summarize overall shifts in microbial diversity by site during giardiasis, we plotted the overall abundance of each identified taxonomic group as a fold change between infected and uninfected samples (Figure 5). In antibiotic treated animals, the taxa with increased diversity in both the foregut and hindgut included the *Rhodocyclaceae* (most notably *Dechloromonas* and *Zoogloea spp*), the *Moraxellaceae*, due to enrichment of *Acinetobacter spp*, the *Flavobacteriales* (*Chryseobacterium* and *Cloacibacterium spp*), and the *Comomonadaceae*, due to the enrichment in unclassified species (Figure 5). Bacterial OTUs that are depleted across all regions of the gut included the *Firmicutes*, and the families *Erysipelotrichaceae* (including *Allobaculum spp*) and *Clostridiaceae* (including *Clostridium spp*). The *Melainabacteria* were enriched in the proximal small intestine, and depleted in the distal small intestine and hindgut. Other taxa (*Lactobacillaceae, Lachnospiraceae, Xanthomonadaceae, Enterobacteriaceae, Pseudomonadaceae, Actinomycetales, Ruminococcaceae, Sphingomonadaceae, Bacteroidales*) were either enriched or depleted in either the foregut or the hindgut. Based on these patterns of diversity during giardiasis, we defined taxa with significant diversity changes across all gut sites as “giardiasis indicator taxa” (underlined groups in Figure 5).

We also summarized alterations in microbial taxa during giardiasis in non-antibiotic treated animals (Supplemental Figure 6). *Rhodocyclaceae* OTUs were enriched in the proximal small intestine, and *Moraxellaceae, Flavobacteriales, Comononadaceae*, and *Bacteroidales* were enriched throughout the small intestine. In the hindgut, only the *Bacteroidales* were enriched. In all other foregut samples, the enriched taxa were depleted in giardiasis. No *Proteobacteria* were found in the hindgut of non-antibiotic-treated animals. *Clostridiacae* were depleted across the intestinal tract of non-antibiotic-treated animals.

Because the bacterial families *Rhodocylaceae* and *Moraxellaceae* were enriched and *Clostridiales* were depleted throughout the course of infection, we created a Murine Giardiasis Dysbiosis Index (MGDI) to summarize shifts in the microbiota during giardiasis. The MGDI combines the relative abundance of *Rhodocylcaceae* and *Moraxellaceae* in the numerator and *Clostridiales* in the denominator, such that a higher MGDI indicates depleted *Clostridiales* and enriched *Rhodocyclaceae* and *Moraxellaceae*. Using this measure, the MGDI steadily rose at all body sites during the duration of infection (Figure 7A). Plotting samples with infection day as an explicit axis shows that elevated GMDI values are only associated with more prolonged infection (Figure 7B).

The MGDI similarly defined microbial shifts associated with infection duration in non-antibiotic-treated animals (Supplemental Figure 7). This shift is magnified in the foregut, yet increased MGDI is also seen in the hindgut. The commensal taxa characterizing the cecum and feces cluster distinctly from the small intestine, in and this in reflected in substantially lower GMDI values in these anatomical regions.

## Discussion

Following ingestion by a mammalian host, *Giardia* cysts transform into motile trophozoites as they pass into the gastrointestinal tract. Trophozoites navigate the lumen of the small intestine, then use the ventral disc to attach to the microvilli of the small intestine, but do not invade the epithelium (11). Attachment to the gut epithelia allows the parasite to resist peristaltic flow and proliferate in this low oxygen, nutrient rich environment. Trophozoites are triggered to transform to cysts, and these eventually exit the host and are disseminated in feces into the environment.

A myriad of factors could influence the pathogenesis of *Giardia*, including host nutritional status, host mucosal immune responses, immune modulation by *Giardia*, variability in the virulence of *Giardia* strains, or even the presence of co-infecting enteropathogens (4). Current hypotheses about how giardiasis induces host symptoms have focused primarily on specific damage induced by parasite attachment to the host gut epithelium or on the production of anti-microbial metabolites by the mammalian host. Parasite colonization within the small intestine occurs in an environment already inhabited by diverse commensal bacteria, yet positive or negative interactions of parasites with the commensal intestinal microbiota that occur during infection have largely been ignored. (4). There is evidence that interactions between *Giardia* and these commensal microbes could contribute to variations in pathogenesis and adverse symptoms associated with giardiasis (47, 54). This idea is supported by evidence of a direct relationship between *Giardia* infection and intestinal bacterial overgrowth (55, 56). Cultures of jejunal juice from patients with active giardiasis revealed increased total bacterial load, increased abundance of *Enterobacteriacae*, and perturbed bile acid homeostasis (56, 57). *Giardia* infection also causes increased bacterial loads mid-infection, and increased bacterial invasiveness post-clearance as a sequela to intestinal barrier disruption (58).

This study is the first high-throughput, culture-independent assessment of the commensal microbiota during *Giardia* infection. Giardiasis perturbs the diversity and abundance of host commensal microbial communities both in the foregut where parasites colonize, and also in the hindgut (Figures 4 and Figure 5, and Supplemental Figures 3-6). This systemic alteration of the murine commensal gut ecology during giardiasis likely contributes to the variation in infection initiation, degree of colonization, induction of pathology in the host, and/or eventual clearance of the parasite (Figure 6) (47, 54).

**Figure 6.**
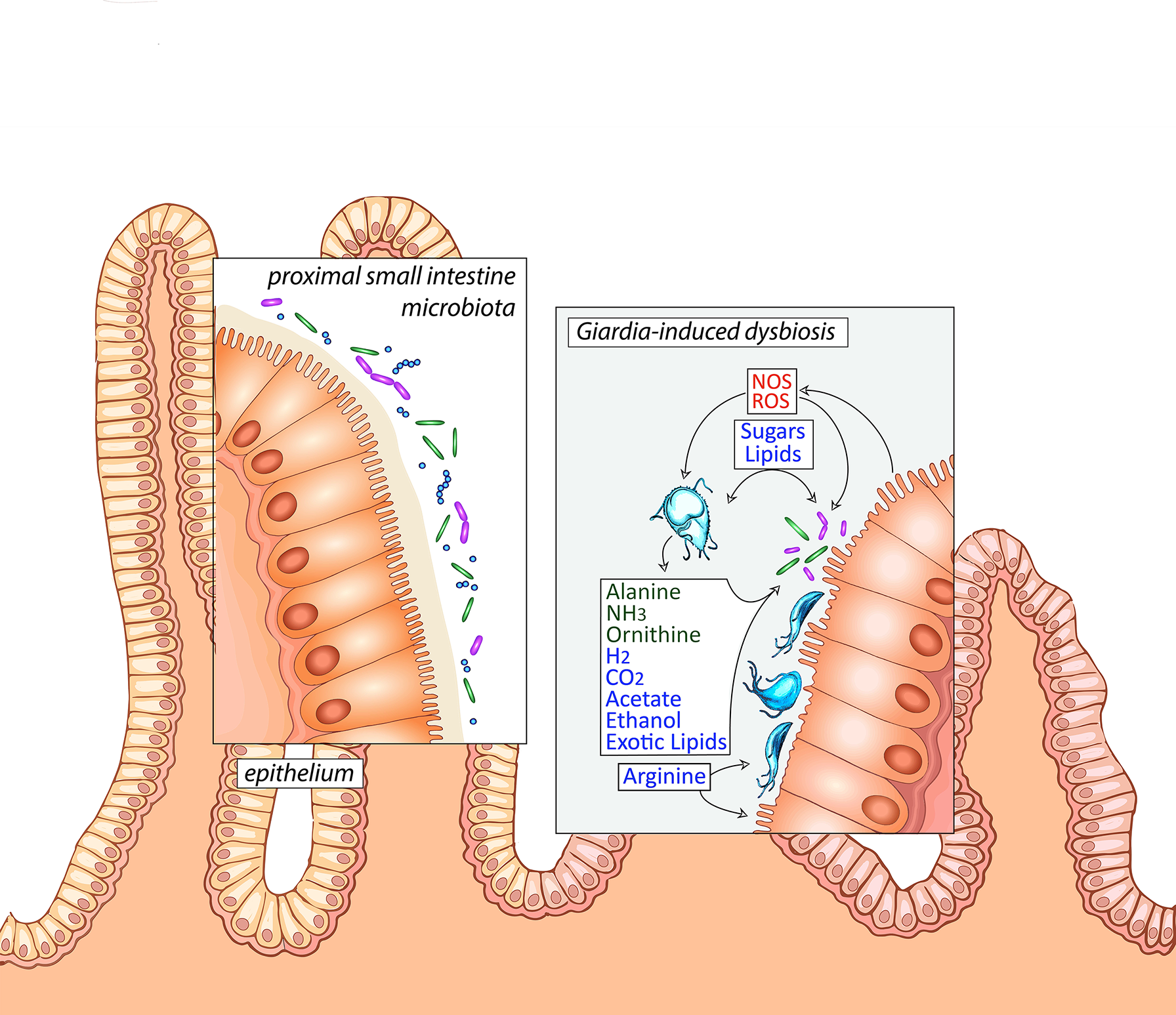
Model of Giardia-induced dysbiosis in the proximal small intestine with implications for disease symptoms. Hypothesized metabolic interactions between fermentative *Giardia* trophozoites with both the host commensal microbiota and host epithelium of the proximal small intestine, the primary site of parasite colonization. Reactive oxygen and nitrogen species (ROS and NOS) are induced by inflammatory responses. Trophozoites ferment sugars and amino acids, and can excrete various waste products depending on the oxygen tension in the surrounding lumen. Primary shifts in diversity of the commensal beta and gamma-*Proteobacteria* and the *Clostridiales* (as well as *Melainabacteria*) are indicated and also summarized in the Giardia Microbial Dysbiosis Index (Figure 7).

### The dysbiosis throughout the murine gastrointestinal tract during giardiasis can be summarized by the “Murine Giardiasis Dysbiosis Index”

Based on our observations, we have developed a Murine Giardiasis Dysbiosis Index (MGDI) as a simplified and quantitative metric of changing community composition during giardiasis (Figure 7 and Supplemental Figure 7). Like other indexes of microbial dysbioses (27), the MGDI summarizes primary ecological shifts in taxon abundance (e.g., the enrichment of *Rhodocylaceae* and *Moraxellaceae* and depletion of *Clostridiales*) and the relative abundance of enriched and depleted microbes. Using this measure, the MGDI steadily increased at all gut regions during the duration of infection, regardless of antibiotic pretreatment (Figure 7 and Supplemental Figure 6). Use of this or similar dysbiosis index, if validated in human samples, could aid in evaluating gut commensal health in infected patients and monitoring for long-term functional gastrointestinal consequences and efficacy of therapies such as fecal microbial transplant (59).

**Figure 7.**
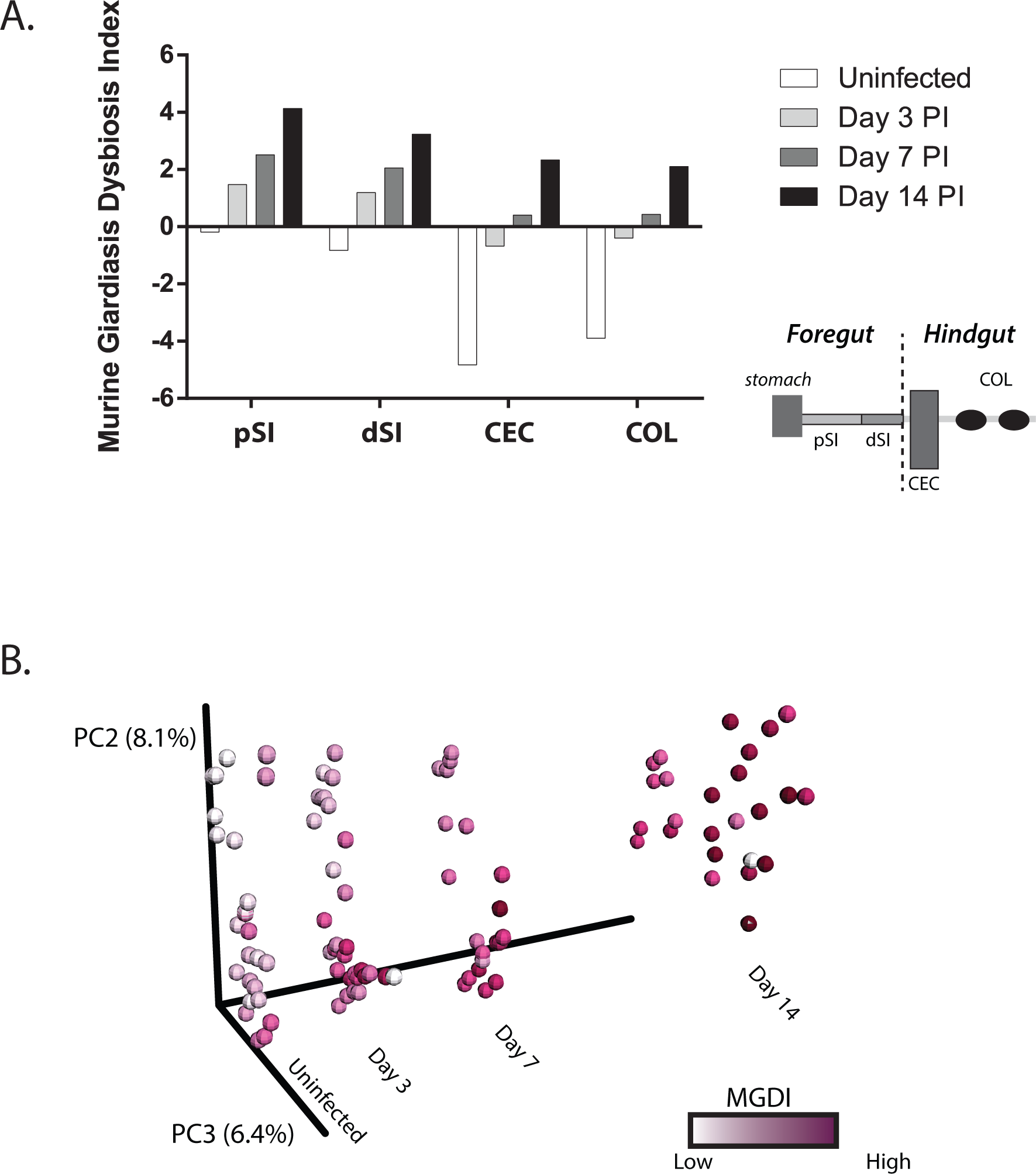
The Murine Giardiasis Dysbiosis Index defines microbial community shifts during the time course of infection in antibiotic-treated animals. The Murine Giardiasis Dysbiosis Index was calculated for each sample by dividing the sum of the abundances of the *Rhodocylaceae* and *Moraxellaceae* families by the abundance of *Clostridiaceae* (see Methods). The average MGDI per body site over the infection time course is presented. The MGDI increases during infection for all body sites analyzed (A). The PCOA (B) displays the infection time course as an explicit axis. Samples are shaded according to their calculated MGDI, with darker shading intensity representing higher MGDI values.

### Giardiasis symptoms may be associated with the ecological shifts induced by parasite metabolism

Parasite colonization of the gut likely alters the local gut chemistry and metabolite availability, changing the gut ecology. In this way, *Giardia* acts an ecosystem engineer (60), modulating the availability of resources to other species in the same environment, and excreting novel waste products such as alanine (61) or exotic lipids (62) that can be metabolized by commensal bacteria. Giardiasis symptoms may reflect the parasite’s impact on the ecology of the gut rather than its direct effect on the host epithelium. By looking at community composition changes during giardiasis and extrapolating to known metabolic capacities of enriched and depleted taxa, we show that *Giardia* infection enriches for metabolically flexible taxa known to thrive with increased oxygen tension, lipid availability, and competition for arginine.

While *Giardia* is commonly considered to be anaerobic due to the lack of many detoxification mechanisms to survive higher oxygen concentrations, trophozoites are, in fact, microaerophilic. Trophozoites are capable of thriving in minimally-oxygenated environments using diaphorase (NADH oxidase) to maintain cellular redox balance (63). Trophozoite glucose fermentation yields several primary fermentation products: acetate, ethanol, alanine, carbon dioxide, and hydrogen (61, 64-69). The end product balance of parasite fermentation is sensitive to local oxygen tension and glucose levels, with highest yield NAD+ or ATP yield at the higher end of tolerated oxygen concentrations (65-68). Though they cannot tolerate oxygen levels that are excessively elevated, trophozoites actively consume oxygen (70, 71) and reap some energetic benefit (61, 67, 72).

Even in the absence of histopathologic evidence of inflammation, giardiasis is associated with redox perturbations (15). Increased diversity and abundance of facultative anaerobes, such as various members of the gamma and beta *Proteobacteria*, can accompany the induction of inflammatory states, and as such, their growth is a sensitive indicator of redox potential in the gastrointestinal environment (73). Alterations in luminal redox potential during giardiasis could be linked to lower levels of anaerobic metabolism by the host microbiota (Figure 6). *Giardia* is proposed to reduce epithelial cell inflammatory responses through decreased host nitric oxide synthesis and reductions in IL-8 (14, 74). *Giardia*-mediated dysbiosis may be caused indirectly by micro-inflammation in the gastrointestinal environment, as has been noted with other opportunistic gut pathogens (22, 27, 75). Inflammation throughout the gut is associated with an overall increased oxygen tension in this generally low oxygen environment, however, and we suggest that alterations in the redox potential during giardiasis are linked to markedly lower levels of anaerobic metabolism by the host microbiota (Figure 6).

The increased abundance of facultative aerobes and strict aerobes we observed during infection could be explained by low-level inflammation associated with giardiasis (76). However, the gamma-*Proteobacteria* taxa enriched in the proximal and distal small intestine during giardiasis are not members of the *Enterobacteriaceae* that are often noted to bloom during gut inflammation (25, 77). While the *Enterobacteriaceae* decreased in diversity, the diversity of the strictly aerobic *Moraxellaceae* and the beta ‐ proteobacteria *Rhodocyclaceae* and *Comomonadaceae* increased. These taxa are generally metabolically flexible with respect to respiratory potential and carbon/nitrogen metabolism, and often metabolize complex carbon polymers including hydrocarbons and lipids (78). Concomitantly, giardiasis-mediated inflammation in the gut resulted in decreased diversity and abundance of obligate anaerobes such as the *Firmicutes* (e.g., *Lactobacillaceae*, *Eryipelotichaeae*, *Ruminococcus*, and *Clostridia*) (27). We also saw decreases in the obligately anaerobic *Melainabacteria* – members of the Cyanobacteria that do not perform photosynthesis and obtain energy by fermentation, producing hydrogen as a waste product. In the human gut, they also synthesize some B and K vitamins, implying that commensal *Melainabacteria* are beneficial to the mammalian host (79, 80).

Compared to commonly studied model eukaryotes, *Giardia* lacks mitochondria and cytochrome mediated oxidative phosphorylation, relying upon limited fermentation for substrate level phosphorylation. Trophozoites also lack *de novo* lipid, purine and pyrimidine synthesis pathways, relying solely on salvage pathways to obtain these nutrients from the local gut environment and microbiota (81-83). Alterations in the overall redox state of the gut would also affect *Giardia* metabolism, perturbing the carbon and nitrogen balance of the local milieu. In turn, these changes could directly influence the metabolic capabilities and diversity of the gut microbiota.

A common symptom of giardiasis is fat malabsorption, with markedly increased lipid concentrations in luminal contents and feces (11, 84, 85). In addition to suffering from characteristic steatorrhea, patients excreting *Giardia* cysts are dyslipidemic, with altered lipid profiles in circulating blood as well as the gut lumen (86). *Giardia* directly perturbs lipid metabolism within the host by scavenging the gut lumen for lipids and by excreting novel end products of lipid metabolism (78); it also contributes to shifts in the local gut microbiota (1) by both excreting novel lipids and influencing the bioavailability of bile acids (62, 84, 87, 88). Bile acids are key endocrine signaling molecules, and perturbation of bile acid homeostasis resulting from giardial metabolism may alter energy metabolism, hepatic health and host cell proliferation (89, 90).

Novel dietary lipids have been shown to notably change microbial community makeup (91). The microbial metabolism of giardial lipids may play an important role in the commensal diversity shifts in the gut. By creating a lipid-rich environment in the gut lumen, *Giardia* colonization may enrich for lipid specialists within the commensal microbiota. The taxa enriched during giardiasis are similar to taxa dominating the microbiome of other hydrocarbon-enriched environments (92-95). *Acinetobacter* spp, which are gamma-*Proteobacteria* within the *Moraxellaceae* family, are notably enriched during giardiasis. *Acinetobacter* species such as *Acinetobacter calcoaceticus* grow using diverse carbon sources, including acetate, ethanol, and exotic hydrocarbons, and have been promoted to degrade these compounds in contaminated environments (96, 97). It is possible that during giardiasis, *Acinetobacter* spp may metabolize these similar lipids as carbon sources. Commensal metabolism of exotic lipids induced during giardiasis could also contribute to this dysbiosis.

Lastly, *Giardia* may also alter the overall gut nitrogen metabolism. Trophozoites also use the arginine dihydrolase pathway (ADiHP) for energy production. This pathway yields ATP via through the conversion of arginine to ammonia and citrulline, with the substrate-level phosphorylation of citrulline yielding ornithine and carbamoylphosphate, resulting in NH_3_, CO_2_ as end products (68). The developmental transition from trophozoites to cysts might also impact nitrogen and carbon metabolism in gut commensal bacteria, as encysting trophozoites slow their catabolism of glucose for energy and switch to using glucose to synthesize N-acetylgalactosamine, a component of the cyst wall. Encysting trophozoites increase arginine catabolism through the ADiHP, possibly to offset energy lost from slowing glycolysis. Both *Giardia* and epithelial cells use arginine for growth, and competition for arginine between the parasite and host could directly influence observed severe malnutrition and increased susceptibility to villus shortening (4, 98). Inducible nitric oxide production by epithelial cells could be impaired by the decreased availability for arginine, and thus alter host cellular immune responses (99).

Thus *Giardia* arginine metabolism could also directly impact commensal microbial metabolism. The enrichment of denitrifiers such as the beta-Proteobacteria (*Rhodocyclaecae*) (Figure 5) that we observe during giardiasis may be related to the metabolism of both vegetative and encysting trophozoites through the excretion of ornithine and ammonium via the arginine dihydrolase pathway. Many of these organisms are noted for their broad metabolic capacities, and some, such as *Zoogloea* and *Dechloromonas* spp. both denitrify and degrade hydrocarbons (100). Other Rhodocyclaecae use organic acids, alcohols, or lipids such as cholesterol for growth (101, 102). Further profiling of metabolic processes of the *Giardia*-infected gut using metabolomics methods could illuminate possible roles of microbial metabolic byproducts during giardiasis.

### Host commensal microbes may limit the initiation or the degree of Giardia colonization

By a general mechanism known as “colonization resistance” (103-105), gut microbiota are thought to antagonize pathogens. Production of bacteriocidins or peroxidases can inhibit or kill pathogenic bacteria. For instance, *Enterococcus*, one of the major constituents of the small intestine, produces superoxide radicals, possibly as a defense mechanism against competitors (106). Understanding how the host gut microbiome limits the initiation of infections could lead to novel therapeutic mechanisms for reducing severity or duration of symptoms, including treatment of at risk children with pre- or probiotics.

Commensal microbiota could limit the initiation or the degree of giardiasis in mice; this may also be reflected in giardiasis in humans. Though convenient for immunologic and genetic manipulations, mice are not a natural host for human strains of *Giardia lamblia*. Reproducible and robust infections of mice with the human-adapted assemblages of *Giardia lamblia* WB (assemblage A) or GS (assemblage B) require pre-treatment of study animals with a cocktail of antibiotics (38). Singer and Young implicated commensal gastrointestinal bacteria in *Giardia’s* pathogenesis when they observed that mice of the same genotype, from two different vendors, were differentially susceptible to infection with strain GS, and that the susceptibility was sensitive to antibiotics and was transferrable by co-housing (47). In fact, host commensal bacteria may have a greater influence over infection susceptibility than the absence of CD4+ T cells (107). More recently, Solaymani-Mohammadi and Singer used a cocktail of antibiotics to successfully infect mice with strain WB (38). *Lactobacillus* probiotic treatment has been reported to reduce the severity and duration of murine giardiasis (108, 109). Symptoms of cramping and steatorrhea do not resolve with antibiotics (110), yet fecal bacteriotherapy has been reported to relieve post-giardial gastrointestinal symptoms for over a year after infection (59).

Direct and indirect parasite-commensal interactions, as well as host-parasite interactions, can be dysregulated during giardiasis through host-mediated inflammatory and microbially-mediated metabolic responses. The specific ecological mechanisms by which *Giardia’s* interactions with commensal microbiota contribute to parasite colonization, infection initiation, pathology, and parasite clearance remain to be explored. A mechanistic understanding of ecological shifts induced during giardiasis could lead to both preventative and therapeutic strategies for treatment. Host-associated microbiota are known to restrict pathogenic colonization (19, 20) and exacerbate or support invasion and pathogenesis (21, 22), and thus could be used therapeutically. While *Lactobacillus* probiotic treatment is reported to reduce the severity and duration of murine giardiasis (108, 109), other probiotic treatments remain unexplored. A clearer understanding of parasite-induced ecological shifts could lead to improved treatments that reduce the severity or duration of symptoms during acute or chronic infection.

## Acknowledgements

This work was supported by to NIH R01AI077571 to SCD and an American Association of Immunologists (AAI) Careers in Immunology Fellowship, NIH AI094492, and NIH AI109591 to SMS. The authors graciously thank Sarah Guest for the schematic representation of the *Giardia*-induced dysbiosis, as well as Kari Hagen, Katherine Karberg, and Chris Nosala for critical reading of the manuscript.

## References

1. Halliez MC, Buret AG. 2013. Extra-intestinal and long term consequences of Giardia duodenalis infections. World J Gastroenterol 19:8974-8985.

2. Al-Mekhlafi HM, Al-Maktari MT, Jani R, Ahmed A, Anuar TS, Moktar N, Mahdy MA, Lim YA, Mahmud R, Surin J. 2013. Burden of Giardia duodenalis infection and its adverse effects on growth of schoolchildren in rural Malaysia. PLoS Negl Trop Dis 7:e2516.

3. Furness BW, Beach MJ, Roberts JM. 2000. Giardiasis surveillance‐‐United States, 1992-1997. Mor Mortal Wkly Rep CDC Surveill Summ 49:1-13.

4. Bartelt LA, Sartor RB. 2015. Advances in understanding Giardia: determinants and mechanisms of chronic sequelae. F1000Prime Rep 7:62.

5. Bartelt LA, Roche J, Kolling G, Bolick D, Noronha F, Naylor C, Hoffman P, Warren C, Singer S, Guerrant R. 2013. Persistent G. lamblia impairs growth in a murine malnutrition model. J Clin Invest 123:2672-2684.

6. Morch K, Hanevik K, Rortveit G, Wensaas KA, Eide GE, Hausken T, Langeland N. 2009. Severity of Giardia infection associated with post-infectious fatigue and abdominal symptoms two years after. BMC Infect Dis 9:206.

7. Morch K, Hanevik K, Rortveit G, Wensaas KA, Langeland N. 2009. High rate of fatigue and abdominal symptoms 2 years after an outbreak of giardiasis. Trans R Soc Trop Med Hyg 103:530-532.

8. Stark D, Barratt JL, van Hal S, Marriott D, Harkness J, Ellis JT. 2009. Clinical significance of enteric protozoa in the immunosuppressed human population. Clin Microbiol Rev 22:634-650.

9. Savioli L, Smith H, Thompson A. 2006. Giardia and Cryptosporidium join the ‘Neglected Diseases Initiative’. Trends Parasitol 22:203-208.

10. Caeiro J-P, DuPont HL. Diarrhea in Adults.

11. Adam RD. 2001. Biology of Giardia lamblia. Clin Microbiol Rev 14:447-475.

12. Ankarklev J, Jerlstrom-Hultqvist J, Ringqvist E, Troell K, Svard SG. 2010. Behind the smile: cell biology and disease mechanisms of Giardia species. Nat Rev Microbiol 8:413-422.

13. Maloney J, Keselman A, Li E, Singer SM. 2015. Macrophages expressing arginase 1 and nitric oxide synthase 2 accumulate in the small intestine during Giardia lamblia infection. Microbes Infect doi: 10.1016/j.micinf.2015.03.006.

14. Stadelmann B, Merino MC, Persson L, Svard SG. 2012. Arginine consumption by the intestinal parasite Giardia intestinalis reduces proliferation of intestinal epithelial cells. PloS one 7:e45325.

15. Cotton JA, Amat CB, Buret AG. 2015. Disruptions of Host Immunity and Inflammation by Giardia Duodenalis: Potential Consequences for Co-Infections in the Gastro-Intestinal Tract. Pathogens 4:764-792.

16. Koot BGP, ten Kate FJW, Juffrie M, Rosalina I, Taminiau JJaM, Benninga Ma. 2009. Does Giardia lamblia cause villous atrophy in children?: A retrospective cohort study of the histological abnormalities in giardiasis. Journal of pediatric gastroenterology and nutrition 49:304-308.

17. Frank DN, Pace NR. 2008. Gastrointestinal microbiology enters the metagenomics era. Current opinion in gastroenterology 24:4-10.

18. McCann KS. 2000. The diversity-stability debate. Nature 405:228-233.

19. Swe PM, Zakrzewski M, Kelly A, Krause L, Fischer K. 2014. Scabies mites alter the skin microbiome and promote growth of opportunistic pathogens in a porcine model. PLoS Negl Trop Dis 8:e2897.

20. Hsiao A, Ahmed AM, Subramanian S, Griffin NW, Drewry LL, Petri WA, Jr., Haque R, Ahmed T, Gordon JI. 2014. Members of the human gut microbiota involved in recovery from Vibrio cholerae infection. Nature 515:423-426.

21. Ferreira RB, Gill N, Willing BP, Antunes LC, Russell SL, Croxen MA, Finlay BB. 2011. The intestinal microbiota plays a role in Salmonella-induced colitis independent of pathogen colonization. PloS one 6:e20338.

22. Winter SE, Thiennimitr P, Winter MG, Butler BP, Huseby DL, Crawford RW, Russell JM, Bevins CL, Adams LG, Tsolis RM, Roth JR, Bäumler AJ. 2010. Gut inflammation provides a respiratory electron acceptor for Salmonella. Nature 467:426-429.

23. Torres MF, Uetanabaro AP, Costa AF, Alves CA, Farias LM, Bambirra EA, Penna FJ, Vieira EC, Nicoli JR. 2000. Influence of bacteria from the duodenal microbiota of patients with symptomatic giardiasis on the pathogenicity of Giardia duodenalis in gnotoxenic mice. J Med Microbiol 49:209-215.

24. Cooks J. 2002. Characterizing ecosystem-level consequences of biological invasions: the role of ecosystem engineers. Oikos 97:153-166.

25. Faber F, Baumler AJ. 2014. The impact of intestinal inflammation on the nutritional environment of the gut microbiota. Immunol Lett 162:48-53.

26. Weingarden A, Gonzalez A, Vazquez-Baeza Y, Weiss S, Humphry G, Berg-Lyons D, Knights D, Unno T, Bobr A, Kang J, Khoruts A, Knight R, Sadowsky MJ. 2015. Dynamic changes in short- and long-term bacterial composition following fecal microbiota transplantation for recurrent Clostridium difficile infection. Microbiome 3:10.

27. Gevers D, Kugathasan S, Denson LA, Vazquez-Baeza Y, Van Treuren W, Ren B, Schwager E, Knights D, Song SJ, Yassour M, Morgan XC, Kostic AD, Luo C, Gonzalez A, McDonald D, Haberman Y, Walters T, Baker S, Rosh J, Stephens M, Heyman M, Markowitz J, Baldassano R, Griffiths A, Sylvester F, Mack D, Kim S, Crandall W, Hyams J, Huttenhower C, Knight R, Xavier RJ. 2014. The treatment-naive microbiome in new-onset Crohn's disease. Cell Host Microbe 15:382-392.

28. Hold GL, Smith M, Grange C, Watt ER, El-Omar EM, Mukhopadhya I. 2014. Role of the gut microbiota in inflammatory bowel disease pathogenesis: what have we learnt in the past 10 years? World J Gastroenterol 20:1192-1210.

29. Molloy MJ, Grainger JR, Bouladoux N, Hand TW, Koo LY, Naik S, Quinones M, Dzutsev AK, Gao JL, Trinchieri G, Murphy PM, Belkaid Y. 2013. Intraluminal containment of commensal outgrowth in the gut during infection-induced dysbiosis. Cell Host Microbe 14:318-328.

30. Benson A, Pifer R, Behrendt CL, Hooper LV, Yarovinsky F. 2009. Gut commensal bacteria direct a protective immune response against Toxoplasma gondii. Cell Host Microbe 6:187-196.

31. Heimesaat MM, Bereswill S, Fischer A, Fuchs D, Struck D, Niebergall J, Jahn HK, Dunay IR, Moter A, Gescher DM, Schumann RR, Gobel UB, Liesenfeld O. 2006. Gram-negative bacteria aggravate murine small intestinal Th1-type immunopathology following oral infection with Toxoplasma gondii. J Immunol 177:8785-8795.

32. Backhed F, Ley RE, Sonnenburg JL, Peterson DA, Gordon JI. 2005. Host-bacterial mutualism in the human intestine. Science 307:1915-1920.

33. Costello EK, Lauber CL, Hamady M, Fierer N, Gordon JI, Knight R. 2009. Bacterial community variation in human body habitats across space and time. Science 326:1694-1697.

34. Dominguez-Bello MG, Costello EK, Contreras M, Magris M, Hidalgo G, Fierer N, Knight R. 2010. Delivery mode shapes the acquisition and structure of the initial microbiota across multiple body habitats in newborns. Proc Natl Acad Sci U S A 107:11971-11975.

35. Zoetendal EG, Heilig HG, Klaassens ES, Booijink CC, Kleerebezem M, Smidt H, de Vos WM. 2006. Isolation of DNA from bacterial samples of the human gastrointestinal tract. Nat Protoc 1:870-873.

36. Caporaso JG, Lauber CL, Walters WA, Berg-Lyons D, Huntley J, Fierer N, Owens SM, Betley J, Fraser L, Bauer M, Gormley N, Gilbert JA, Smith G, Knight R. 2012. Ultra-high-throughput microbial community analysis on the Illumina HiSeq and MiSeq platforms. ISME J 6:1621-1624.

37. Gilbert JA, Meyer F, Jansson J, Gordon J, Pace N, Tiedje J, Ley R, Fierer N, Field D, Kyrpides N, Glockner FO, Klenk HP, Wommack KE, Glass E, Docherty K, Gallery R, Stevens R, Knight R. 2010. The Earth Microbiome Project: Meeting report of the "1 EMP meeting on sample selection and acquisition" at Argonne National Laboratory October 6 2010. Stand Genomic Sci 3:249-253.

38. Solaymani-Mohammadi S, Singer SM. 2011. Host immunity and pathogen strain contribute to intestinal disaccharidase impairment following gut infection. J Immunol 187:3769-3775.

39. Caporaso JG, Kuczynski J, Stombaugh J, Bittinger K, Bushman FD, Costello EK, Fierer N, Pena AG, Goodrich JK, Gordon JI, Huttley GA, Kelley ST, Knights D, Koenig JE, Ley RE, Lozupone CA, McDonald D, Muegge BD, Pirrung M, Reeder J, Sevinsky JR, Turnbaugh PJ, Walters WA, Widmann J, Yatsunenko T, Zaneveld J, Knight R. 2010. QIIME allows analysis of high-throughput community sequencing data. Nat Methods 7:335-336.

40. Love MI, Huber W, Anders S. 2014. Moderated estimation of fold change and dispersion for RNA-seq data with DESeq2. Genome Biol 15:550.

41. Vazquez-Baeza Y, Pirrung M, Gonzalez A, Knight R. 2013. EMPeror: a tool for visualizing high throughput microbial community data. Gigascience 2:16.

42. Lozupone CA, Stombaugh J, Gonzalez A, Ackermann G, Wendel D, Vazquez-Baeza Y, Jansson JK, Gordon JI, Knight R. 2013. Meta-analyses of studies of the human microbiota. Genome Res 23:1704-1714.

43. Wagner Mackenzie B, Waite DW, Taylor MW. 2015. Evaluating variation in human gut microbiota profiles due to DNA extraction method and inter-subject differences. Front Microbiol 6:130.

44. Anders S, Huber W. 2010. Differential expression analysis for sequence count data. Genome Biol 11:R106.

45. Frank DN, Robertson CE, Hamm CM, Kpadeh Z, Zhang T, Chen H, Zhu W, Sartor RB, Boedeker EC, Harpaz N, Pace NR, Li E. 2011. Disease phenotype and genotype are associated with shifts in intestinal associated microbiota in inflammatory bowel diseases. Inflammatory bowel diseases 17:179-184.

46. Morrison TB, Ma Y, Weis JH, Weis JJ. 1999. Rapid and sensitive quantification of Borrelia burgdorferi infected mouse tissues by continuous fluorescent monitoring of PCR. J Clin Microbiol 37:987-992.

47. Singer SM, Nash TE. 2000. The role of normal flora in Giardia lamblia infections in mice. J Infect Dis 181:1510-1512.

48. Solaymani-Mohammadi S, Singer SM. 2013. Regulation of intestinal epithelial cell cytoskeletal remodeling by cellular immunity following gut infection. Mucosal Immunol 6:369-378.

49. Roberts-Thomson IC, Stevens DP, Mahmoud AA, Warren KS. 1976. Giardiasis in the mouse: an animal model. Gastroenterology 71:57-61.

50. Gillin FD, Boucher SE, Reiner DS. 1987. Stimulation of In-Vitro Encystation of Giardia-Lamblia by Small Intestinal Conditions. Clinical Research 35:475A.

51. Lujan HD, Mowatt MR, Byrd LG, Nash TE. 1996. Cholesterol starvation induces differentiation of the intestinal parasite Giardia lamblia. Proc Natl Acad Sci U S A 93:7628-7633.

52. Gillon J, Al Thamery D, Ferguson A. 1982. Features of small intestinal pathology (epithelial cell kinetics, intraepithelial lymphocytes, disaccharidases) in a primary Giardia muris infection. Gut 23:498-506.

53. Peterson DA, Frank DN, Pace NR, Gordon JI. 2008. Metagenomic approaches for defining the pathogenesis of inflammatory bowel diseases. Cell host & microbe 3:417-427.

54. Eckmann L. 2003. Mucosal defences against Giardia. Parasite Immunol 25:259-270.

55. Müller N, von Allmen N. 2005. Recent insights into the mucosal reactions associated with Giardia lamblia infections. International journal for parasitology 35:1339-1347.

56. Tandon BN, Tandon RK, Satpathy BK, Shriniwas. 1977. Mechanism of malabsorption in giardiasis: a study of bacterial flora and bile salt deconjugation in upper jejunum. Gut 18:176-181.

57. Tomkins AM, Wright SG, Drasar BS, James WPT, Unit M, Diseases T. 1978. Bacterial colonization of jejunal mucosa in giardiasis. Transactions of the Royal Society of Tropical Medicine and Hygiene 72.

58. Chen TL, Chen S, Wu HW, Lee TC, Lu YZ, Wu LL, Ni YH, Sun CH, Yu WH, Buret AG, Yu LC. 2013. Persistent gut barrier damage and commensal bacterial influx following eradication of Giardia infection in mice. Gut Pathog 5:26.

59. Morken MH, Valeur J, Norin E, Midtvedt T, Nysaeter G, Berstad A. 2009. Antibiotic or bacterial therapy in post-giardiasis irritable bowel syndrome. Scandinavian journal of gastroenterology 44:1296-1303.

60. Jones C, Lawton J, Shachak M. 1994. Organisms as Ecosystem Engineers. Oikos 69:373-386.

61. Paget TA, Raynor MH, Shipp DW, Lloyd D. 1990. Giardia lamblia produces alanine anaerobically but not in the presence of oxygen. Mol Biochem Parasitol 42:63-67.

62. Yichoy M, Duarte TT, De Chatterjee A, Mendez TL, Aguilera KY, Roy D, Roychowdhury S, Aley SB, Das S. 2011. Lipid metabolism in Giardia: a post-genomic perspective. Parasitology 138:267-278.

63. Lloyd D. 2004. 'Anaerobic protists': some misconceptions and confusions. Microbiology 150:1115-1116.

64. Lindmark DG. 1980. Energy metabolism of the anaerobic protozoan Giardia lamblia. Molecular and Biochemical Parasitology 1:1-12.

65. Schofield PJ, Costello M, Edwards MR, O'Sullivan WJ. 1990. The arginine dihydrolase pathway is present in Giardia intestinalis. Int J Parasitol 20:697-699.

66. Schofield PJ, Edwards MR, Kranz P. 1991. Glucose metabolism in Giardia intestinalis. Mol Biochem Parasitol 45:39-47.

67. Edwards MR, Schofield PJ, O’Sullivan WJ, Costello M. 1992. Arginine metabolism during culture of Giardia intestinalis. Mol Biochem Parasitol 53:97-103.

68. Schofield PJ, Edwards MR, Matthews J, Wilson JR. 1992. The pathway of arginine catabolism in Giardia intestinalis. Mol Biochem Parasitol 51:29-36.

69. Lloyd D, Harris JC, Maroulis S, Wadley R, Ralphs JR, Hann AC, Turner MP, Edwards MR. 2002. The "primitive" microaerophile Giardia intestinalis (syn. lamblia, duodenalis) has specialized membranes with electron transport and membrane-potential-generating functions. Microbiology 148:1349-1354.

70. Paget TA, Manning P, Jarroll EL. 1993. Oxygen uptake in cysts and trophozoites of Giardia lamblia. J Eukaryot Microbiol 40:246-250.

71. Paget TA, Kelly ML, Jarroll EL, Lindmark DG, Lloyd D. 1993. The effects of oxygen on fermentation in Giardia lamblia. Mol Biochem Parasitol 57:65-71.

72. Adam RD. 2001. Biology of Giardia lamblia. Clinical microbiology reviews 14.

73. Rigottier-Gois L. 2013. Dysbiosis in inflammatory bowel diseases: the oxygen hypothesis. ISME J 7:1256-1261.

74. Mastronicola D, Falabella M, Forte E, Testa F, Sarti P, Giuffre A. 2015. Antioxidant defence systems in the protozoan pathogen Giardia intestinalis. Mol Biochem Parasitol doi: 10.1016/j.molbiopara.2015.12.002.

75. Lupp C, Robertson ML, Wickham ME, Sekirov I, Champion OL, Gaynor EC, Finlay BB. 2007. Host mediated inflammation disrupts the intestinal microbiota and promotes the overgrowth of Enterobacteriaceae. Cell Host Microbe 2:119-129.

76. Albenberg L, Esipova TV, Judge CP, Bittinger K, Chen J, Laughlin A, Grunberg S, Baldassano RN, Lewis JD, Li H, Thom SR, Bushman FD, Vinogradov SA, Wu GD. 2014. Correlation between intraluminal oxygen gradient and radial partitioning of intestinal microbiota. Gastroenterology 147:1055-1063 e1058.

77. Winter SE, Baumler AJ. 2014. Dysbiosis in the inflamed intestine: chance favors the prepared microbe. Gut microbes 5:71-73.

78. Mendez TL, De Chatterjee A, Duarte T, De Leon J, Robles-Martinez L, Das S. 2015. Sphingolipids, Lipid Rafts, and Giardial Encystation: The Show Must Go On. Curr Trop Med Rep 2:136-143.

79. Wrighton KC, Castelle CJ, Wilkins MJ, Hug LA, Sharon I, Thomas BC, Handley KM, Mullin SW, Nicora CD, Singh A, Lipton MS, Long PE, Williams KH, Banfield JF. 2014. Metabolic interdependencies between phylogenetically novel fermenters and respiratory organisms in an unconfined aquifer. ISME J 8:1452-1463.

80. Di Rienzi SC, Sharon I, Wrighton KC, Koren O, Hug LA, Thomas BC, Goodrich JK, Bell JT, Spector TD, Banfield JF, Ley RE. 2013. The human gut and groundwater harbor non-photosynthetic bacteria belonging to a new candidate phylum sibling to Cyanobacteria. Elife 2:e01102.

81. Jarroll EL, Muller PJ, Meyer EA, Morse SA. 1981. Lipid and carbohydrate metabolism of Giardia lamblia. Mol Biochem Parasitol 2:187-196.

82. Aldritt SM, Tien P, Wang CC. 1985. Pyrimidine salvage in Giardia lamblia. J Exp Med 161:437-445.

83. Wang CC, Aldritt S. 1983. Purine salvage networks in Giardia lamblia. J Exp Med 158:1703-1712.

84. Das S, Stevens T, Castillo C, Villasenor A, Arredondo H, Reddy K. 2002. Lipid metabolism in mucous dwelling amitochondriate protozoa. Int J Parasitol 32:655-675.

85. Katelaris P, Seow F, Ngu M. 1991. The effect of Giardia lamblia trophozoites on lipolysis in vitro. Parasitology 103 Pt 1:35-39.

86. Bansal D, Bhatti HS, Sehgal R. 2005. Altered lipid parameters in patients infected with Entamoeba histolytica, Entamoeba dispar and Giardia lamblia. Br J Biomed Sci 62:63-65.

87. Das S, Schteingart CD, Hofmann AF, Reiner DS, Aley SB, Gillin FD. 1997. Giardia lamblia: evidence for carrier-mediated uptake and release of conjugated bile acids. Exp Parasitol 87:133-141.

88. Gillin FD, Gault MJ, Hofmann AF, Gurantz D, Sauch JF. 1986. Biliary lipids support serum-free growth of Giardia lamblia. Infect Immun 53:641-645.

89. Houten SM, Watanabe M, Auwerx J. 2006. Endocrine functions of bile acids. EMBO J 25:1419-1425.

90. Keitel V, Kubitz R, Haussinger D. 2008. Endocrine and paracrine role of bile acids. World J Gastroenterol 14:5620-5629.

91. Marques TM, Wall R, O'Sullivan O, Fitzgerald GF, Shanahan F, Quigley EM, Cotter PD, Cryan JF, Dinan TG, Ross RP, Stanton C. 2015. Dietary trans-10, cis-12-conjugated linoleic acid alters fatty acid metabolism and microbiota composition in mice. Br J Nutr 113:728-738.

92. Abbai NS, Pillay B. 2013. Analysis of hydrocarbon-contaminated groundwater metagenomes as revealed by high-throughput sequencing. Mol Biotechnol 54:900-912.

93. Paisio CE, Quevedo MR, Talano MA, Gonzalez PS, Agostini E. 2014. Application of two bacterial strains for wastewater bioremediation and assessment of phenolics biodegradation. Environ Technol 35:1802-1810.

94. Chen Y, Li C, Zhou Z, Wen J, You X, Mao Y, Lu C, Huo G, Jia X. 2014. Enhanced biodegradation of alkane hydrocarbons and crude oil by mixed strains and bacterial community analysis. Appl Biochem Biotechnol 172:3433-3447.

95. Singleton DR, Dickey AN, Scholl EH, Wright FA, Aitken MD. 2015. Complete Genome Sequence of a Novel Bacterium within the Family Rhodocyclaceae That Degrades Polycyclic Aromatic Hydrocarbons. Genome Announc 3.

96. Abdel-El-Haleem D. 2003. Acinetobacter: environmental and biotechnological applications. African Journal of Biotechnology 2:71-74.

97. Khoramnia A, Ebrahimpour A, Beh BK, Lai OM. 2011. Production of a solvent, detergent, and thermotolerant lipase by a newly isolated Acinetobacter sp. in submerged and solid-state fermentations. J Biomed Biotechnol 2011:702179.

98. Ventura LL, Oliveira DR, Viana JC, Santos JF, Caliari MV, Gomes MA. 2013. Impact of protein malnutrition on histological parameters of experimentally infected animals with Giardia lamblia. Exp Parasitol 133:391-395.

99. Eckmann L, Laurent F, Langford TD, Hetsko ML, Smith JR, Kagnoff MF, Gillin FD. 2000. Nitric oxide production by human intestinal epithelial cells and competition for arginine as potential determinants of host defense against the lumen-dwelling pathogen Giardia lamblia. J Immunol 164:1478-1487.

100. Farkas M, Tancsics A, Kriszt B, Benedek T, Toth EM, Keki Z, Veres PG, Szoboszlay S. 2015. Zoogloea oleivorans sp. nov., a floc-forming, petroleum hydrocarbon-degrading bacterium isolated from biofilm. Int J Syst Evol Microbiol 65:274-279.

101. Smalley NE, Taipale S, Marco P, Doronina NV, Kyrpides N, Shapiro N, Woyke T, Kalyuzhnaya MG. 2015. Functional and genomic diversity of methylotrophic Rhodocyclaceae: description of the new species Methyloversatilis discipulorum sp. nov. Int J Syst Evol Microbiol doi: 10.1099/ijs.0.000190.

102. Chiang YR, Ismail W, Heintz D, Schaeffer C, Van Dorsselaer A, Fuchs G. 2008. Study of anoxic and oxic cholesterol metabolism by Sterolibacterium denitrificans. J Bacteriol 190:905-914.

103. Sharma A, Dhayal D, Singh OP, Adak T, Bhatnagar RK. 2013. Gut microbes influence fitness and malaria transmission potential of Asian malaria vector Anopheles stephensi. Acta Trop 128:41-47.

104. Flavia Nardy A, Freire-de-Lima CG, Morrot A. 2015. Immune Evasion Strategies of Trypanosoma cruzi. J Immunol Res 2015:178947.

105. Mooney JP, Lokken KL, Byndloss MX, George MD, Velazquez EM, Faber F, Butler BP, Walker GT, Ali MM, Potts R, Tiffany C, Ahmer BM, Luckhart S, Tsolis RM. 2015. Inflammation-associated alterations to the intestinal microbiota reduce colonization resistance against non-typhoidal Salmonella during concurrent malaria parasite infection. Sci Rep 5:14603.

106. Wang X, Huycke MM. 2007. Extracellular superoxide production by Enterococcus faecalis promotes chromosomal instability in mammalian cells. Gastroenterology 132:551-561.

107. Singer SM, Nash TE. 2000. T-cell-dependent control of acute Giardia lamblia infections in mice. Infect Immun 68:170-175.

108. Goyal N, Tiwari RP, Shukla G. 2011. Lactobacillus rhamnosus GG as an Effective Probiotic for Murine Giardiasis. Interdisciplinary perspectives on infectious diseases 2011:795219.

109. Shukla G, Sidhu RK. 2011. Lactobacillus casei as a probiotic in malnourished Giardia lamblia-infected mice: a biochemical and histopathological study. Canadian Journal of Microbiology 135:127-135.

110. Wright SG, Tomkins AM, Ridley DS. 1977. Giardiasis: clinical and therapeutic aspects. Gut 18:343-350.

